# Spontaneous regression of micro-metastases following primary tumor excision: a critical role for primary tumor secretome

**DOI:** 10.1101/2020.03.12.986992

**Authors:** Lee Shaashua, Anabel Eckerling, Boaz Israeli, Gali Yanovich, Ella Rosenne, Suzana Fichman-Horn, Ido Ben Zvi, Liat Sorski, Ronit Satchi-Fainaro, Tamar Geiger, Erica K. Sloan, Shamgar Ben-Eliyahu

## Abstract

Numerous case studies have reported spontaneous regression of recognized metastases following primary tumor (PT) excision, but underlying mechanisms are elusive. Here we present a model of metastases regression and latency following PT excision, and identify potential underlying mechanisms. Using MDA-MB-231^HM^ human breast cancer cells that express highly sensitive luciferase, we were able to monitor early stages of spontaneous metastases development in BALB/c nu/nu mice. Removal of the PT caused marked regression of the smallest micro-metastases, but not of larger metastases, and *in vivo* supplementation of tumor secretome diminished this regression, suggesting that PT-secreted factors promote early metastatic growth. Correspondingly, cancer cell conditioned medium reduced apoptosis and enhanced MDA-MB-231^HM^ adhesion *in vitro*. To identify specific mediating factors, cytokine array and proteomic analysis of MDA-MB-231^HM^ secretome were conducted. Results identified significant enrichment of angiogenesis, growth factors binding and activity, focal adhesion, metalloprotease regulation, and apoptosis regulation processes. Simultaneous *in vivo* blockade of four secreted key potential mediators of these processes, IL-8, PDGFaa, Serpin E1 (PAI-1), and MIF, arrested development of micro-metastases in the presence of the PT. Interestingly, using the public TCGA provisional dataset, high protein levels of these four factors were correlated with poor survival in a cohort of lung adenocarcinoma patients. These results demonstrate regression and latency of micro-metastases following PT excision, and a crucial role for PT-secretome in promoting early metastatic stages in MDA-MB-231^HM^ xenografts. If generalized, such findings can suggest novel approaches to control minimal residual disease during and following PT excision.

## Introduction

Surgical removal of the primary tumor is a cornerstone of cancer treatment. Nevertheless, this life-saving approach has been suggested to accelerate postoperative progression of minimal residual disease through processes triggered by the surgical procedure itself [1, 2], or by elimination of inhibitory signaling from the primary tumor [3].

However, various case studies report the opposite effect; spontaneous postoperative regression of evident cancer metastasis [4–7]. Such postoperative regression, although rare, has been clearly documented in most types of cancer, including breast cancer [6, 7], and has been most commonly reported for lung metastases [8–13].

Several underlying mechanisms have been suggested to elicit spontaneous postoperative regression of residual malignant foci [10, 14, 15], including surgical trauma [5], and elimination of stimulating factors secreted by the primary tumor or induced by its presence [7, 15]. Unfortunately, postulated mechanisms have not been empirically tested, as no animal model of spontaneous regression exists. The need to study potential interactions between the primary tumor (PT) and metastases is stressed by the current realization that the presence of a primary tumor is a systemic disease, and that a continuous crosstalk between the PT, its microenvironment, and distant organs, plays a significant role in disease etiology and progression [16, 17].

Herein, for the first time, we present an animal model of spontaneous regression of metastases following PT removal. This model employs a highly sensitive luciferase reporter of cancer cells, which enables the study of early-stage micro-metastases. We found that development of micro-metastases is supported by the numerous factors secreted from the PT, and that removal of the PT and its secreted factors induces regression of early-stage metastases. Specific potential factors were identified, and an *in vivo* neutralization of four of them in the presence of the PT halted progression of metastases.

## Methods

### Cancer model

A highly metastatic variant of the triple-negative breast adenocarcinoma cell line, MDA-MB-231^HM^ (a gift from Dr. Zhou Ou, Fudan University, Shanghai Cancer Center, China) was transduced with a codon-optimized firefly luciferase-mCherry vector as previously described [18]. Cells were cultured in Dulbecco’s modified Eagle’s medium (DMEM; Thermo Fisher Scientific) supplemented with 10% fetal bovine serum (FBS), 1% glutamax (Thermo Fisher Scientific), 4.5g/L D-glucose and 110mg/L sodium pyruvate. Cells were maintained at 37 °C and 5% CO2, and were mycoplasma-free.

### *In vivo* model of spontaneous metastasis

Mice were housed under SPF conditions on 12 h dark/light cycle. Eight-week old female BALB/c nu/nu mice (University of Adelaide, Australia or Envigo, Israel) were injected with 2×10^5^ cells in 20 μl PBS into the fourth mammary fat pad (under 2% isoflurane anesthesia) to form a primary tumor. Primary tumors were measured by caliper, and volume was calculated by the formula: (length × width^2^)×0.5. Metastases were assessed by bioluminescence imaging using an IVIS spectrum apparatus (Perkin Elmer) following i.p. injection of 150mg/kg D-luciferin sodium salt (Regis Technologies). Once metastatic foci reached a total count of 10^6^ photons/s (~ 3-4 weeks post injection), the primary tumor (average size of 80mm^3^) was excised, and complete removal was verified by bioluminescence imaging. Follow up of metastatic progression was conducted by bioluminescence imaging. At the end point, animals were euthanized 10 min following luciferin injection, and lungs and lymph nodes were harvested for *ex vivo* imaging. All procedures were approved by Monash University or Tel-Aviv University Animal Ethics Committees.

### Surgical procedures

Primary tumor resection was performed under anesthesia with 2% isoflurane. A small incision in the skin was performed without injuring the peritoneal cavity to excise the PT. Following a complete removal of the PT (which was verified by bioluminescence imaging), the skin was immediately sutured. For the sham surgery, an identical incision was made, however the PT was untouched. The lesion was immediately sutured.

### Tumor cell- conditioned medium (CM) preparation and use

CM was produced by incubating ~80% confluent MDA-MB-231^HM^ cells with serum-free medium for 24 h at 37°C, 5% CO2. SM (i.e. serum-free medium, SM) used as control. For *in vivo* supplementation of tumor secretome, CM was collected from ~30 million cells and filtered through 0.45μm strainer. CM (and the same volume of SM as control) was then concentrated by 3 kDa Amicon filters (Merck-Millipore) to a final volume of 100 μl/mouse and stored at −20°C until usage. On implantation day, Alzet osmotic minipumps (model 1003D) were loaded with either CM or SM and implanted i.p. immediately following tumor excision. Animals were randomly assigned to the experimental conditions. Based on an estimation that an excised primary tumor contains ~50×10^6^ secretome-producing cells (in ~80 mm^3^ primary tumor), we estimate that the amount of secretome factors released/day by the osmotic minipumps used is ~1/8 than the amount produced *in vivo* /day by the primary tumor.

### Apoptosis and adhesion studies

Cancer cells were cultured in growth media, washed and seeded in either SM or CM. For adhesion studies, cells were imaged for 8 hours, in 20 min intervals, using the IncuCyte system (Essen Bioscience). The number of adhered cells was determined. For apoptosis studies, cells were incubated in CM or SM for 24 hrs, then washed and stained for AnnexinV-FITC (R&D systems) and 7AAD (R&D systems). Using flow cytometry, percent of live/dead/early apoptotic cells was determined.

### Tube formation assay

24-well plates wells were layered with 50μl basement membrane extracellular matrix (Cultrex BME; Trevigen), and incubated for 30 min at 37°C, 5% CO2. Then, human umbilical vein endothelial cells (HUVEC), reconstituted in either CM or SM, were seeded. Cells were imaged at 20 min intervals for 24 h using IncuCyte system (Essen Bioscience). Average cells area (μm^2^) was analyzed by the IncuCyte software and the number of tubes was counted.

### Human cytokine analysis

Human Cytokine Array (R&D systems, Proteome Profiler™ Array, ARY022) was used to compare relative expression levels of 102 soluble human proteins in CM samples, and in plasma samples from (i) mice bearing a primary tumor, (ii) mice one day following primary tumor excision, and (iii) control mice. Each plasma sample was pooled from 3 mice. Analysis and quantification were used by Protein Array Analyzer in Image J software.

### Mass spectrometry-based secretome analysis

CM samples were centrifuged to eliminate cell debris followed by filtration and concentration using 3 kDa Amicon filters. Concentrated medium was mixed at a 1:1 ratio with 8M urea and filtered again to reach a final volume of ~100ul. Prior to protein digestion, proteins from the filtered samples were incubated with 1mM dithiothreitol followed by 5mM iodoacetamide. Proteins were digested overnight with LysC/Trypsin mix (Promega) and sequencing-grade modified Trypsin (Promega) at room temperature, followed by desalting and concentration on C_18_ StageTips [19]. Prior to MS analysis, peptides were eluted from StageTips using 80% acetonitrile, vacuum-concentrated and diluted in MS loading buffer (2% acetonitrile, 0.1% formic acid). Liquid Chromatography-Mass Spectrometry (LC-MS) was performed using the nano-ultra high performance liquid chromatography system (UHPLC) (Easy-nLC1000, Thermo Fisher Scientific), followed by MS analysis on the Q-Exactive Plus mass spectrometer (Thermo Fisher Scientific). Peptides were separated by reverse-phase chromatography (50 cm long EASY-Spray PepMap columns; Thermo Fisher Scientific) with a 140 min linear gradient of water/acetonitrile. MS analysis was performed using a top10 method in which every high resolution MS scan was followed by fragmentation of the 10 most abundant peaks by higher-energy collisional dissociation (HCD).

### Proteomics data analysis

Mass spectrometry (MS) raw files were analyzed by MaxQuant [20], and the Label-free quantification algorithm [21]. MS/MS spectra were referenced to the Uniprot human proteome by the Andromeda search engine. A false discovery rate (FDR) of 0.01 was used on both the peptide and protein levels based on a decoy database. Statistical analysis was performed using Perseus software [22], String database (www.string-db.org) and Cytoscape software. Enrichment analysis was performed relative to the identified secretome using Gene Ontology annotations from UniProt (Fisher exact test with an FDR threshold of 0.02).

### Neutralizing antibodies and their *in vivo* use

Mouse anti-human monoclonal antibodies (R&D systems) were used to neutralize CXCL8/IL-8 (IgG1 Clone # 6217), Serpin E1/PAI-1 (IgG1 Clone # 242816), PDGF-AA (IgG1 Clone # 114506), and MIF (IgG1 Clone # 12302), and monoclonal mouse IgG1 served as isotype control (IgG1 Clone # 11711). All antibodies/isotype control were injected once a day (1μg each/mouse), for two days.

### Histology

To identify lung micro-metastases, lungs were imaged *ex vivo*, and lobes with bioluminescence-verified micro-metastases were collected, fixed in 4% paraformaldehyde, and paraffin embedded. Lung sections (6 μm) were stained for Hematoxylin and Eosin (H&E). Metastatic foci were confirmed by a pathologist.

### Patient outcome analysis

Using the public TCGA provisional dataset (cBioPortal) [23, 24], we studied patient outcome in a lung adenocarcinoma cohort, stratifying for patients with altered protein levels of IL-8, PDGFaa, Serpin E1 and MIF, based on relative protein levels in RPPA assay. Survival curve was conducted according to Kaplan Meier Analysis, and a hazard ratio was calculated in the R software.

### Statistical analysis

Repeated-measures or factorial analysis of variance (ANOVA), with a pre-determined significance level of 0.05 were conducted. Provided significant group differences were found, Fisher’s protected least significant differences (Fisher’s PLSD) contrasts were performed to test pair-wise post hoc comparisons. Paired or unpaired student’s t-test was performed for comparing two experimental conditions (following F-test of equality of variance). All statistical tests were two sided. Throughout this study, technical replicates, if conducted, were averaged and used to reliably represent a biological replicate.

## Results

To examine the effect of primary tumor excision on metastatic growth, MDA-MB-231^HM^ cells were injected into the mammary fat pad of nude mice to form a primary tumor. In this orthotopic model, distant metastases are formed spontaneously in lymph nodes and lungs 2-4 weeks following tumor implantation. When chest-localized bioluminescent signal of metastases reached a total flux of 10^6^ photons/s, primary tumors were excised. A dramatic decrease in metastatic signal, of up to 100-fold, was evident in mice along the 24 h following tumor excision, p<0.0001 (Fig. 1A–B) (but not immediately following excision). These metastases have regressed into latent foci, as (i) no increase in *in vivo* bioluminescent signal occurred for 50 days post-excision (Fig. 1B–C), and as (ii) microscopic malignant foci were evident on both the day after tumor excision (day 1) and 50 days later, as confirmed by H&E staining of lungs and lymph nodes (Fig. 1D). To test whether regression is specific to early metastatic stages, tumors were excised from mice bearing either small (10^6^ photons/sec) or large (10^7^ photons/sec) metastases; regression was significantly less prominent in larger metastases, p<0.000 (Fig. 1E).

**Figure 1:**
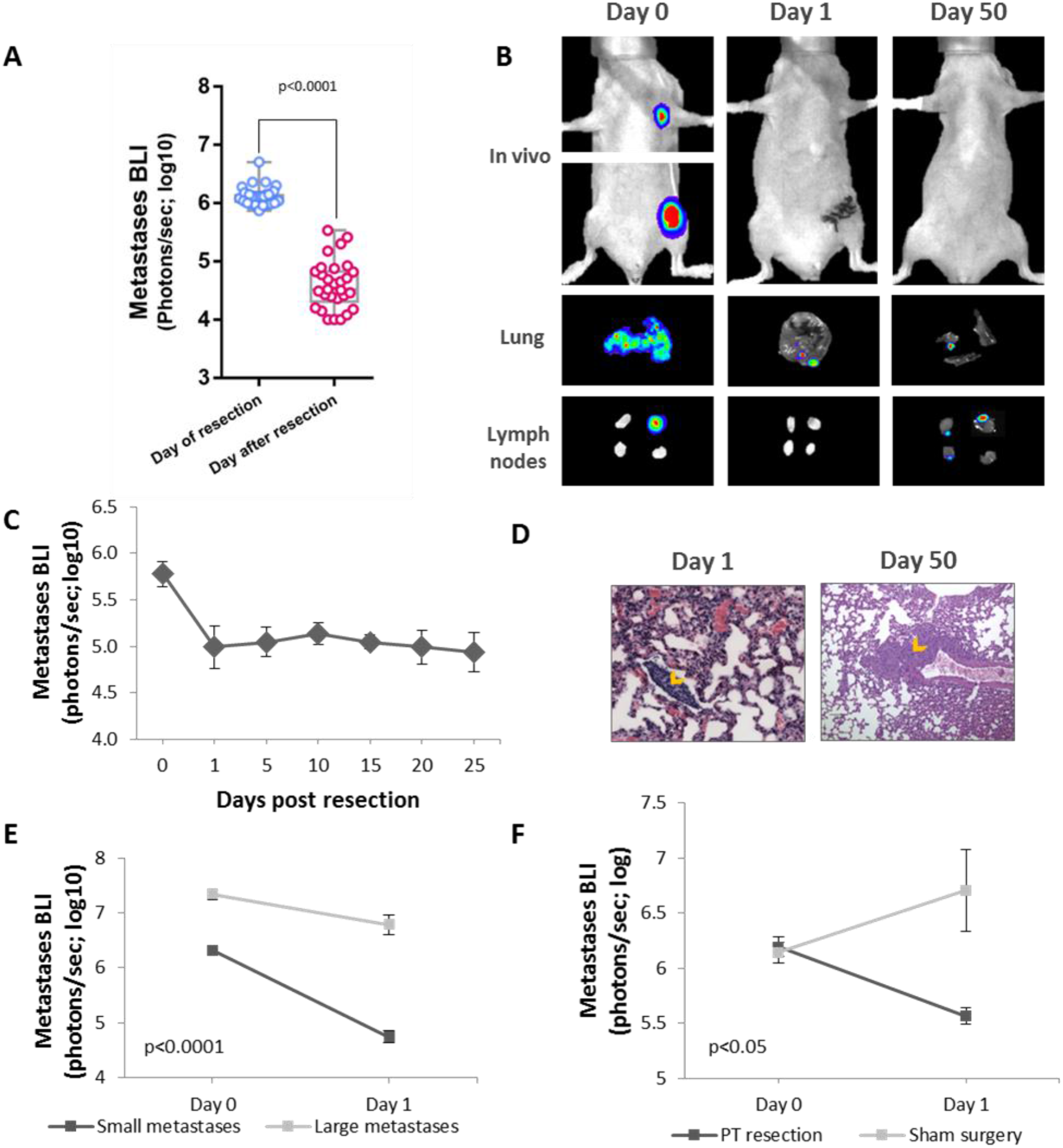
Excision of the primary tumor elicits regression of early-stage metastases. **(A)** *In vivo* quantification of lung and lymph node metastasis by bioluminescence imaging (BLI) immediately before (day 0), and after primary tumor (PT) resection (day 1) (n=29). Whiskers represent min and max points. **(B)** Representative images of lungs and lymph node (LN) metastases, *in vivo* and ex-vivo at day 0, and post-operative days 1 and 50, following tumor excision. **(C)** *In vivo* BLI of metastases over time (n=6). **(D)** Representative images of H&E staining of lung sections at post-operative days 1 and 50. Orange arrows indicate micro-metastases. **(E)** *In vivo* quantification of lung and LN metastasis by BLI before (day 0), and after PT resection (day 1) in mice bearing small (n=7) or large (n=9) metastases. **(F)** *In vivo* quantification of lung and LN metastasis by BLI before (day 0), and after (day 1) sham surgery or PT resection. Error bars in 1C,1E, 1F represent mean ± SE.

As several mechanisms have been proposed to explain spontaneous regression, we first sought to distinguish between the effects elicited by the surgical procedure itself (such as inflammation, and vascular insufficiency) and those that are related to the removal of the tumor mass, including the elimination of primary tumor-derived pro-metastatic secreted factors (i.e., growth factors, cytokines, and angiogenic factors). To this end, mice underwent either sham surgery (sparing the primary tumor) or primary tumor excision. Metastases regressed significantly following excision of the primary tumor, while continued to increase in mice subjected to sham surgery, p<0.05 (Fig. 1F).

As the spontaneous regression was elicited by the removal of the primary tumor, we hypothesized that the primary tumor secretome supports the survival and growth of distant micro-metastases, and thus its elimination by surgery results in metastatic regression. As regression was more prominent in small metastatic foci (Fig. 1E), we investigated events that characterize the early stages of metastasis development. We found that medium conditioned by MDA-MB-231^HM^ tumor cells, containing tumor-derived secreted factors (compared to serum-free medium), enhanced viability, p<0.001 (Fig. 2A) and adhesion, p<0.01 (Fig. 2B) of cancer cells, and induced tube formation of human endothelial cells, p<0.0001 (Fig. 2C). To study the *in vivo* effect of secreted factors on metastasis, osmotic mini-pumps containing tumor-cell conditioned medium (vs. serum-free medium) were implanted at the time of tumor excision to partly replenish the primary tumor secretome. Mice that were treated with tumor-cell conditioned medium showed 10-fold less regression than mice that received serum-free medium, p<0.005 (Fig. 2D), indicating that secreted factors from the primary tumor support survival of distant metastases. We estimate that the amount of secreted factors released daily by the osmotic mini-pumps was ~⅛ of the amount produced daily *in vivo* by the primary tumor (see Methods).

**Figure 2:**
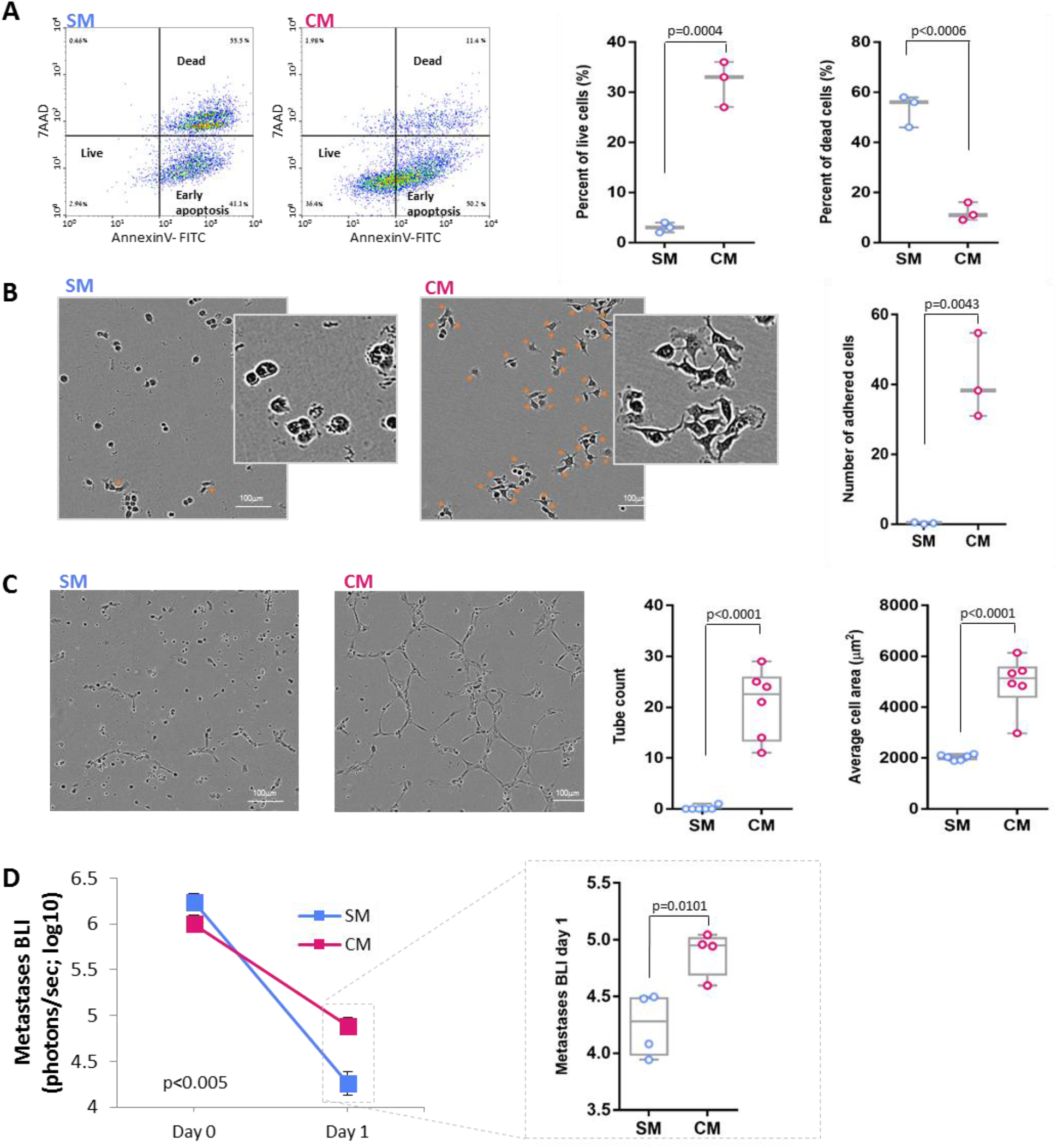
CM effects on pro-metastatic processes and metastasis. **(A)** Representative images and quantification of flow cytometry for AnnexinV and 7AAD of MDA-MB-231^HM^ cells that were grown in serum-free medium (SM) or MDA-MB-231^HM^ tumor-cell conditioned medium (CM). **(B)** Representative images and quantification of adhered cancer cells incubated with SM or CM. Orange arrows mark adhered cells; Scale bar - 100 μm. **(C)** Representative images and quantification of tube formation by human endothelial cells incubated with SM or CM on a layer of basement membrane extracellular matrix; Scale bar - 100 μm. **(D)** *In vivo* quantification of lung and LN metastases by BLI before (day 0), and after PT resection (day 1) in mice that received SM or CM, simultaneously with tumor excision, (n=4 per group). Whiskers in Fig. 2A–2D represent min and max points. Error bars in Fig. 2D represent mean ± SE.

To identify factors whose elimination mediate the spontaneous regression of metastases following primary tumor excision, tumor secretome was analyzed employing a human cytokine array. We analyzed 4 conditions: (i) tumor cell-conditioned medium (CM); and plasma samples from (ii) non-tumor bearing mice, (iii) primary tumor-bearing mice, and (iv) mice one day following tumor excision. We identified 28 cytokines that were highly expressed in cancer cell CM. We then excluded factors that were also evident in plasma of non-tumor bearing mice and/or in plasma of mice following tumor excision (see Table S1). This selection method pointed at four factors that are known to exert pro-metastatic or pro-survival activities, and thus their removal may induce spontaneous regression: interleukin-8 (IL-8), platelet-derived-growth-factor-aa (PDGFaa), serpin E1 (also known as plasminogen-activator-inhibitor-1; PAI-1), and macrophage-migration-inhibitory-factor (MIF). The presence of these 4 factors in CM was validated using ELISA (not shown).

To complement this “narrow down” approach, we used an unbiased mass spectrometry proteomic analysis of CM and identified 2,600 proteins, 359 of which were annotated as extracellular factors by Gene Ontology (GO) analysis [25], including the four factors identified using the cytokine array. Pathway enrichment analysis of the 359 extracellular proteins identified proteins engaged in key steps of metastasis including apoptosis, angiogenesis, growth factor activity, focal adhesion, and metalloenzyme regulation. Examination of protein-protein interaction networks using the String database identified pathways involved in the early stages of metastasis (Fig. 3A)[26].

**Figure 3:**
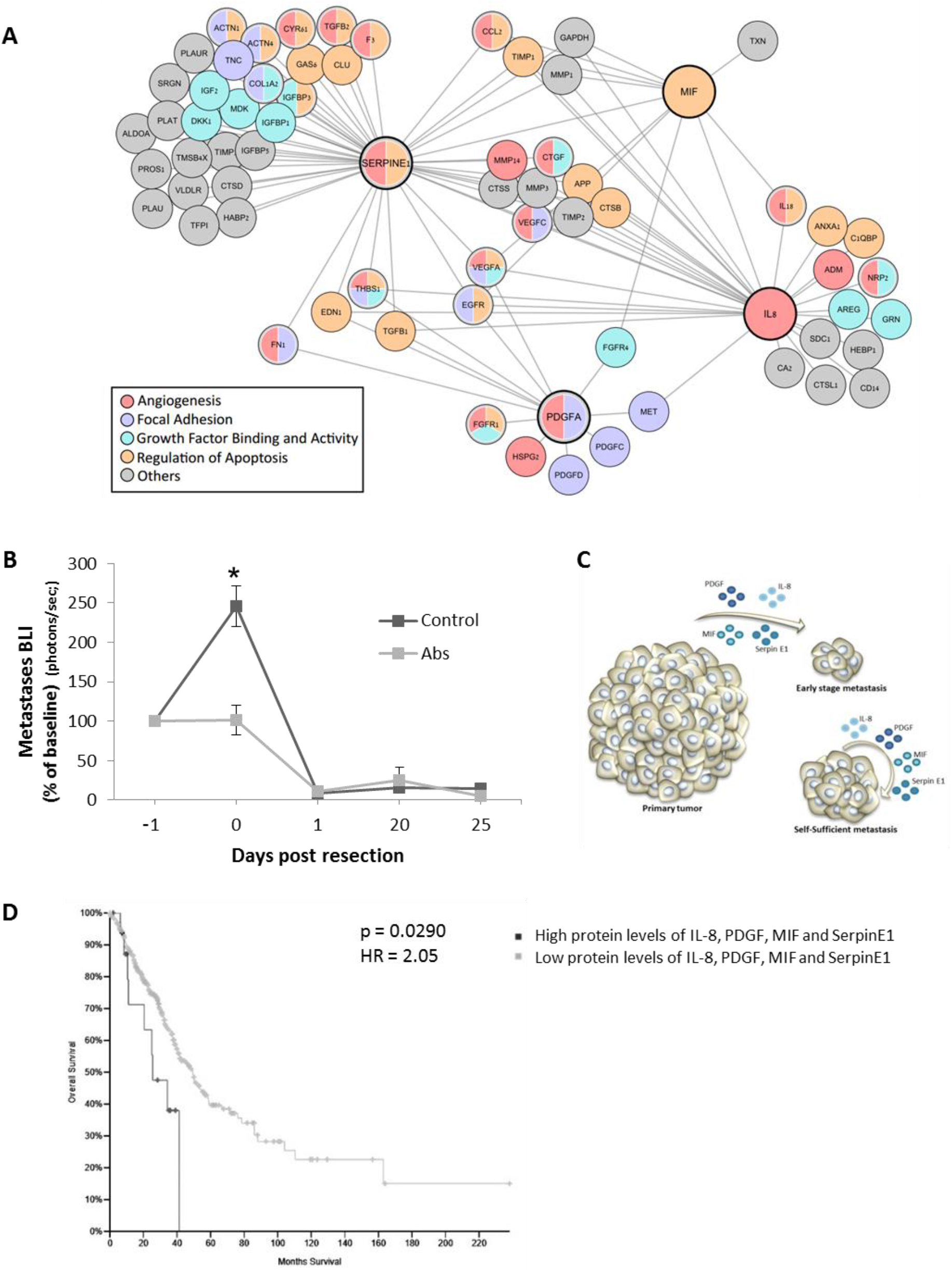
Secreted factors and pathways potentially underlying metastatic regression. **(A)** Proteomic and GO enrichment analysis identified a network of proteins and enriched pathways. Proteins in this scheme are those connected to at least one of the 4 key factors (denoted by a thick line). **(B)** *In vivo* long-term quantification of metastasis following cocktail administration of four neutralizing antibodies (IL-8, MIF, PDGF-aa, and serpin E1; n=4) or IgG control (n=5). Data represent mean ± SE. **(C)** A scheme of the hypothesized model based on our results, suggesting that primary tumor secretome is crucial for survival of early stage micro-metastases, but not for larger metastases. **(D)** Kaplan-Meier analysis of patients stratified by high/low proteins levels of IL-8, MIF, PDGF-aa, and serpin E1 (n=18 in high protein levels, and n= 338 in low protein levels). Antibodies – Abs.

We hypothesized that if these four factors are among those that drive metastatic growth, their *in vivo* blockade would partly mimic the effect of primary tumor removal and inhibit metastasis. To investigate this, a cocktail of mouse anti-human antibodies for IL-8, PDGFaa, serpin E1 and MIF, versus IgG control, were injected 24 h prior to tumor excision, at an early metastatic stage (chest bioluminescence < 10^6^ p/s). Antibody neutralization of these factors completely blocked metastatic progression, p<0.0001 (Fig. 3B). As could be expected, no difference in regression of metastases was evident between these two groups on post-excision day-1, likely as excision completely removed all secreted factors in both groups. These findings suggest that these primary tumor-secreted factors are among the factors crucial for survival of early stage micro-metastases, while established larger metastatic foci may be self-sufficient (Fig. 3C). To explore the clinical relevance of these tumor-derived factors, we retrospectively studied outcome in a cohort of lung adenocarcinoma patients from the public TCGA provisional dataset. Kaplan Meier Analysis revealed that high protein levels of IL-8, PDGFaa, Serpin E1 and MIF were associated with significantly lower survival with a hazard ratio of 2.05, p<0.05 (Fig. 3D).

## Discussion

The findings of this study indicate a crucial supportive role for MDA-MB-231^HM^ tumor secretome in survival and progression of distant metastatic foci, and demonstrate the phenomena of metastatic regression following primary MDA-MB-231^HM^ tumor removal. We identified four tumor-secreted proteins, IL-8, MIF, SerpinE1, and PDGFaa, which are known to maintain and promote metastases, and showed that their in vivo neutralization in the presence of the PT arrested metastatic progression. Using the public TCGA provisional dataset, high protein levels of these four factors were correlated with poor survival in a cohort of lung adenocarcinoma patients. Our findings suggest that tumor-secreted factors may act through multiple mechanisms to affect both malignant cells and/or their microenvironment (Fig. 2). The findings provide an opportunity to empirically study metastatic regression into latent foci, and to explore mediating mechanisms, with the goal of identifying novel prophylactic and therapeutic strategies.

It is well established that the primary tumor secretes numerous factors (e.g., growth factors, angiogenic factors, cytokines, exosomes, and hormones) that promote its growth, modulate its microenvironment [16], and induce a “pre-metastatic niche” at distant organs [17, 27]. Nonetheless, the presence of a primary tumor is generally believed to be primarily metastasis-inhibitory [28], through other mechanisms. The current study shows that primary tumor secretome can be a vital factor in promoting metastasis during early stages of metastatic formation. Additionally, this phenomenon might also occur in substantial portion of patients or in other animal models, but are overlooked due to low detection sensitivity of early micrometastases. Specifically, the use of high-sensitive codon-optimized luciferase-2 (luc-2), combined with the early time of metastatic development studied herein, enabled us to detect metastases and their regression into latent foci in these early stages, which presumably would not occur at more advanced metastatic stages or not detected under lower sensitivity. Indeed, herein larger micrometastases showed lower or no regression, potentially given their ability to secrete sufficient amount of the necessary factors.

Several mechanisms were previously suggested to underlie spontaneous regression. As in this study surgical procedure without primary tumor excision did not elicit any regression of metastases, the potential mechanism of surgical trauma and its associated processes is negated in the current setting. Interestingly, several studies reported spontaneous regression of metastases following radiation or cryotreatment of the primary tumor [29, 30]. These cases could be explained by the current results, as these treatments probably cease the secretion of many factors by the manipulated primary tumor.

Our work proposes a role for IL-8, MIF, SerpinE1, and PDGFaa in maintaining and promoting micrometastases, through shared and distinct pathways. All four factors have been shown to promote angiogenesis and/or to suppress tumor apoptosis. Specifically, IL-8 is a pro-angiogenic and pro-inflammatory chemokine that was shown to (i) promote epithelial to mesenchymal transition (EMT) and invasion of tumor cells [31–35], and (ii) enhance malignant survival. SerpinE1 (PAI-1) was shown to promote metastases by: (i) increasing thrombosis which support angiogenesis [36–38], (ii) inducing pro-survival and anti-apoptotic activities in both epithelial and tumor cells [39–41], and (iii) enhancing tumor invasion [42–44]. The proangiogenic factor, PDGFaa, was shown to act as a survival factor, to inhibit apoptosis, to increase vessel density [45], and to stimulate reorganization of actin [46]. The pro-inflammatory factor, MIF, was shown to promote metastases by (i) initiating the NF-ĸB signaling cascade resulting in the secretion of pro-inflammatory cytokines such as IL-8, TNF-α, IL-1, and IL-6, (ii) promoting MMPs activity, (iii) increasing tumor-infiltration of myeloid-derived suppressor cells [47–48], (iv) promoting EMT [49], and (v) exerting pro-survival [50] and anti-apoptotic activities [51]. Overall, these PT secreted factors are likely to promote metastatic progression in micrometastatic niches, especially when autocrine release of growth supporting factors is insufficient.

Given the multiple factors potentially mediating the pro-metastatic effects of MDA-MB-231^HM^ secretome, we herein tested only the combined effect of blocking four key secreted factors known to promote malignant progression. Other secreted factors are likely to also be involved in supporting metastatic progression. Different tumors, specifically syngeneic lines, should be similarly studied to test the generalizability of the novel phenomenon observed in the current study, and would potentially identify other or common mediating factors.

The results of this study suggest that the perioperative period could be exploited to control minimal residual disease by blocking primary tumor support for micro-metastases, and by targeting pro-metastatic factors released by minimal residual disease, both before and after surgery. Specifically, identifying critical pro-metastatic factors secreted by the primary tumor in each patient, based on malignant tissue biopsy before surgery, may enable perioperative use of individually tailored combination of specific neutralizing antibodies. Such a treatment can also counteract the metastasis-promoting effects of surgery [52,1] and tilt the balance toward eradication of metastases or arrest of their growth. The perioperative period harbor numerous risk factors for metastatic progression, and was thus suggested to present a window of opportunity to exert high impact on long-term cancer outcomes through various interventions [1,53]. The herein findings may explain the rare but validated clinical phenomenon of spontaneous regression of metastases following PT excision. Analyses of serum samples before and after PT resection may be used, if available, to suggest specific mediating factors. Last, it is worth to note that the prevalence of post-operative regression of micro-metastases in cancer patients is currently unknown due to technical limitations, and that such a process may occur in a substantial portion of patients.

## Acknowledgments

We would like to thank Sabine Albold, Hagar Lavon, Noam Cohen and Galia Tiram for their technical assistance, and for Dr. Steven Cole and Dr. Ruth Scherz-Shouval for their evaluation of the manuscript.

## Author contributions

LS, SBE, and ES conceived, designed, and wrote the manuscript. LS, BI, IBZ, ER and SFH collected or analyzed the data. TG and GY analyzed and interpreted the proteomic data. AE, LS and RSF took part in conceiving experiments as well as in manuscript revision.

## Data and materials availability

The proteomics data have been deposited to the ProteomeXchange Consortium (accession code: PXD008384; Username; Password: WJ1e4ARO). All other data is available in the main text or the supplementary materials.

