## Supplementary material for "Spontaneous regression of micro-metastases following primary tumor excision: a critical role for primary tumor secretome": Table S1

Supplementary Materials

### **Table S1 – Cytokines emerging from the cytokine array**

| **CM *in vitro* secretion** | **Plasma levels of tumor bearing mice** | **Ratio of plasma levels from tumor-bearing/control mice** | **Ratio of plasma levels from tumor- bearing/excised tumor mice** |
|---|---|---|---|
| Serpin E1 | + | + | + |
| PDGF-AA | + | + | + |
| IL-8 | + | + | + |
| MIF | + | + | + |
| Dkk-1 | + | + | + |
| MCP-1 | + | + | |
| Thrombospondin-1 | + | + | |
| uPAR | + | + | |
| IL-22 | + | + | |
| Pentraxin-3 | + | | |
| FGF-19 | + | | |
| IL-17A | + | | |
| M-CSF | + | | |
| IL-11 | + | | |
| IL-6 | + | | |
| Osteopontin | + | | |
| Angiopoietin-2 | + | | |
| Vitamin D | + | | |
| SDF-1 | + | | |
| FGF | + | | |
| LIF | | | |
| Angiogenin | | | |
| EMMPRIN | | | |
| GDF-15 | | | |
| VEGF | | | |
| IGFBP-3 | | | |
| Cystatin C | | | |
| GM-CSF | | | |

**Table S1:** List of cytokines emerging from the cytokine array, and criteria used for the selection process to suggest potential prominent factors. The left column represents the 28 upregulated cytokines in MDA-MB-231$^{HM}$ *in vitro* CM. Out of these, 20 cytokines were upregulated in plasma of tumor-bearing mice. Then we selected for cytokines which their levels in tumor bearing mice were upregulated compared to both control mice and mice following tumor resection
